# Morphogenesis and differentiation of embryonic vascular smooth muscle cells in zebrafish

**DOI:** 10.1101/414003

**Authors:** Thomas R. Whitesell, Paul Chrystal, Jae-Ryeon Ryu, Nicole Munsie, Ann Grosse, Curtis French, Matthew L. Workentine, Rui Li, Lihua Julie Zhu, Andrew Waskiewicz, Ordan J. Lehmann, Nathan D. Lawson, Sarah J. Childs

## Abstract

Despite the critical role of vascular mural cells (smooth muscle cells and pericytes) in supporting the endothelium of blood vessels, we know little of their early morphogenesis and differentiation. *foxc1b:EGFP* expressing cells in zebrafish associate with the vascular endothelium (*kdrl*) and co-express a smooth muscle marker (*acta2*), but not a pericyte marker (*pdgfrβ*). The expression of *foxc1b* in early peri-endothelial mesenchymal cells allows us to follow the morphogenesis of mesenchyme into *acta2* expressing vascular smooth muscle cells. We show that mural cells expressing different markers associate with vessels of different diameters, depending on their embryonic location and developmental timing, suggesting marker expression is predictive of functional differences. We identify gene expression signatures for an enriched vascular smooth muscle cell population (*foxc1b + acta2)* and all smooth muscle *(acta2)* using fluorescence-activated cell sorting and RNA-Seq. Finally, we demonstrate that progressive loss of *foxc1a*/*foxc1b* results in decreased smooth muscle cell coverage. Together, our data highlight the early cellular dynamics and transcriptome profiles of smooth muscle cells *in vivo,* using *foxc1b* as a unique tool to probe vascular smooth muscle cell differentiation.

**Summary Statement:** Tracing the morphogenesis and transcriptome of early vascular smooth muscle cells using foxc1b

## Introduction

Smooth muscle cells and pericytes (collectively known as vascular mural cells) encircle and stabilize the underlying endothelium, deposit extracellular matrix proteins, and provide contractility to blood vessels; however, they have distinct morphologies (Lindahl *et al*., 1997; Santoro, Pesce and Stainier, 2009; Stratman *et al*., 2009, 2017; Armulik, Genové and Betsholtz, 2011; Carmeliet and Jain, 2011; Whitesell *et al*., 2014). Vascular smooth muscle cells typically form a multilayer, continuous sheath on large caliber blood vessels subject to higher blood pressure. Found outside the basement membrane in mammals, vascular smooth muscle cells provide support and contractility to the underlying endothelium (Etchevers *et al*., 2001; Santoro, Pesce and Stainier, 2009; Olson and Soriano, 2011; Whitesell *et al*., 2014; Stratman *et al*., 2017). Pericytes are typically punctate, non-continuous cells with individual processes that wrap around smaller diameter blood vessels creating peg and socket connections with the underlying endothelium. Pericytes are essential in regulating vascular permeability, for example, in the blood-brain barrier, and are more prevalent at branching points of blood vessels, turbulent areas of blood flow, or bends in complex blood vessels (Hellström *et al*., 1999; Etchevers *et al*., 2001; Bergers and Song, 2005; Winkler, Bell and Zlokovic, 2010; Armulik, Genové and Betsholtz, 2011; Olson and Soriano, 2011; Whitesell *et al*., 2014; Van Dijk *et al*., 2015; Craggs *et al*., 2015; Trost *et al*., 2016; He *et al*., 2016; Stratman *et al*., 2017; Birbrair *et al*., 2017; Dias Moura Prazeres *et al*., 2017; Berthiaume *et al*., 2018). Highlighting their importance, defects in vascular mural cells lead to cerebrovascular dysfunction including cerebral small vessel disease and stroke (Joutel *et al*., 1996; Craggs *et al*., 2014; French *et al*., 2014; Baron-Menguy *et al*., 2017). For example, loss of vascular smooth muscle cells results in thoracic aneurysm and dissections (TAAD) (Guo *et al*., 2009) and increased vessel size due to poor smooth muscle contraction (Abrams *et al*., 2016), while loss of pericytes leads to increased brain permeability, aneurysm, and hemorrhage (Lindahl *et al*., 1997; Moura, Lemos and Oliveira, 2017). Recent morphological and transcriptomic characterization in mouse suggests there are sub-types of pericytes and smooth muscle cells that vary in gene expression and morphology depending on their position in the vascular tree (Hartmann *et al*., 2015; Grant *et al*., 2017; Vanlandewijck *et al*., 2018). Further illustrating their complexity, mural cell subtypes are typically distinguished by morphology, location, contractile function, and molecular markers (Hill *et al*., 2015; He *et al*., 2016; Birbrair *et al*., 2017; Dias Moura Prazeres *et al*., 2017). In addition, there are multiple developmental origins of mural cells, as vascular smooth muscle cells and pericytes of the head arise from neural crest and mesoderm, while trunk vascular smooth muscle cells have a mesodermal origin (Etchevers *et al*., 2001; Pouget, Pottin and Jaffredo, 2008; Wang *et al*., 2014; Cavanaugh, Huang and Chen, 2015).

Smooth muscle actin (*acta2/αsma*) and transgelin (*tagln/sm22α-b*) are the most widely used vascular smooth muscle markers (Solway *et al*., 1995; Hellström *et al*., 1999; Schildmeyer *et al*., 2000; Georgijevic *et al*., 2007; Santoro, Pesce and Stainier, 2009; Seiler, Abrams and Pack, 2010; Whitesell *et al*., 2014; Stratman *et al*., 2017). Pericytes on the other hand, are typically labelled with platelet-derived growth factor receptor-beta (*pdgfrβ*) (Armulik, Genové and Betsholtz, 2011; Ando *et al*., 2016; He *et al*., 2016; Trost *et al*., 2016; Grant *et al*., 2017; Stratman *et al*., 2017; Berthiaume *et al*., 2018) or *ng2/cspg4* (Armulik, Genové and Betsholtz, 2011; He *et al*., 2016; Trost *et al*., 2016; Underly *et al*., 2017; Berthiaume *et al*., 2018), in addition to *tbx18* (Birbrair *et al*., 2017; Guimar??es-Camboa *et al*., 2017), *notch3* (Wang *et al*., 2014; Henshall *et al*., 2015; Tao *et al*., 2017), *nestin* (Dias Moura Prazeres *et al*., 2017), *rgs5* (Armulik, Genové and Betsholtz, 2011; He *et al*., 2016), *abcc9* (He *et al*., 2016; Vanlandewijck *et al*., 2018), and *desmin* (Georgijevic *et al*., 2007; Armulik, Genové and Betsholtz, 2011; Trost *et al*., 2016). A key caveat is that even classic markers for smooth muscle cells (*acta2, tagln*) and pericytes (*pdgfrβ, ng2/cspg4*), do not label every vascular mural cell (Santoro, Pesce and Stainier, 2009; Whitesell *et al*., 2014; Hartmann *et al*., 2015; He *et al*., 2016; Birbrair *et al*., 2017; Dias Moura Prazeres *et al*., 2017; Grant *et al*., 2017). Mural cell heterogeneity varies with location within an organism, cellular lineage, function, phenotype, and morphology (Georgijevic *et al*., 2007; Santoro, Pesce and Stainier, 2009; Armulik, Genové and Betsholtz, 2011; He *et al*., 2016; Dias Moura Prazeres *et al*., 2017), and consequently, heterogeneity in marker expression makes it difficult to identify a single marker for consistent labelling of vascular mural cells. In addition, there are a lack of smooth muscle markers during early development, particularly in zebrafish (Georgijevic *et al*., 2007), and the common pericyte markers have been shown to be expressed in numerous other cell types (He *et al*., 2016; Ivanova and Orekhov, 2016).

Cerebral pericytes are labelled by *pdgfrβ* expression as early as 48 hpf (hours post fertilization) in zebrafish embryos (Wang *et al*., 2014), while *ng2/cspg4* is expressed by 72 hpf (Wang *et al*., 2014). In zebrafish, *acta2* and *tagln* transgenic expression is first detectable around 3 days post fertilization (dpf), with robust vascular smooth muscle expression by 4 dpf (Georgijevic *et al*., 2007; Majesky, 2007; Santoro, Pesce and Stainier, 2009; Seiler, Abrams and Pack, 2010; Whitesell *et al*., 2014; Gays *et al*., 2017). Using transmission electron microscopy, we previously observed mural cells with a mesenchymal phenotype associated with the endothelium as early as 48 hpf, and showed close association with endothelium correlating with vascular stabilization (Liu *et al*., 2007; Lamont *et al*., 2010). However, the earliest stages of vascular mural cell development in zebrafish cannot be observed using current transgenic markers.

Since many vascular smooth muscle cells of the head are neural crest-derived, we focused on markers that label vascular mural cells early in development and are expressed in neural crest, such as the forkhead box domain transcription factor *foxc1b*. Loss of *Foxc1* in mouse leads to severe defects in head mesenchyme, branchial arch and anterior somite development (Kume *et al*., 1998, 2001). In humans, it is also associated with anterior segment dysgenesis in the eye, and more recently, also with cerebral small vessel disease (Mears *et al*., 1998; Nishimura *et al*., 1998; French *et al*., 2014; Seo *et al*., 2017; Avasarala, Jones and Rogers, 2018). Mouse models show that *Foxc1* mutants have disrupted mural cell contacts on the endothelium, and cerebral hemorrhage (Kume *et al*., 1998; Siegenthaler *et al*., 2013; French *et al*., 2014; Prasitsak *et al*., 2015; Mishra *et al*., 2016). *Foxc1/c2* play important roles in numerous processes including lymphatic development (Seo *et al*., 2006; Van Steensel *et al*., 2009; Sasman *et al*., 2012; Fatima *et al*., 2016); vascular development (Kume *et al*., 2001; Yamagishi *et al*., 2003; Seo *et al*., 2006, 2012; De Val *et al*., 2008; Hayashi and Kume, 2008a, 2008b; Skarie and Link, 2009; Veldman and Lin, 2012; Koo and Kume, 2013; Prasitsak *et al*., 2015; Mishra *et al*., 2016); cardiac outflow tract development (Seo and Kume, 2006; Kodo *et al*., 2017); somitogenesis (Kume *et al*., 2001; J. M. Topczewska *et al*., 2001; Tamimi *et al*., 2006; Qiu *et al*., 2016); neural crest and skeletal development (Seo and Kume, 2006; Inman *et al*., 2013; Koo and Kume, 2013; Kodo *et al*., 2017; Seo *et al*., 2017); and hematopoiesis (Omatsu *et al*., 2014; Jang *et al*., 2015), and diseases such as glaucoma and Axenfeld-Rieger syndrome (Kume *et al*., 2001; Tamimi *et al*., 2006; Berry *et al*., 2008; Skarie and Link, 2009; French *et al*., 2014; Mishra *et al*., 2016; Souzeau *et al*., 2017). In mouse, *Foxc1/c2* are expressed in neural crest (Kume *et al*., 1998; Seo *et al*., 2012, 2017; Siegenthaler *et al*., 2013), and in the endothelium (Kume *et al*., 1998, 2001; Koo and Kume, 2013). *Foxc1* is also expressed in brain pericytes and smooth muscle cells and is co-expressed with *Pdgfrβ* and *Acta2* as early as E14.5 (Zarbalis *et al*., 2007; Siegenthaler *et al*., 2013; Prasitsak *et al*., 2015; Mishra *et al*., 2016). In zebrafish, there are two foxc1 paralogs, *foxc1a* and *foxc1b* (J. M. Topczewska *et al*., 2001; Skarie and Link, 2009; Chen *et al*., 2017). *foxc1b* is first expressed during the involution of mesendoderm in early gastrulation, and by 33 hpf, *foxc1b* is expressed in the pharyngeal arch mesenchyme (Jolanta M. Topczewska *et al*., 2001). Knockdown of *foxc1a/b* in zebrafish shows a reduction in *acta2* positive smooth muscle cells in the ventral aorta and aortic arch arteries (French *et al*., 2014), suggesting a role in mural cell development. *Foxc1* expression is also reported in the endothelium of mouse and in the trunk endothelium of zebrafish (Kume *et al*., 1998; J. M. Topczewska *et al*., 2001; Skarie and Link, 2009; Siegenthaler *et al*., 2013; Prasitsak *et al*., 2015; Mishra *et al*., 2016; Winkler *et al*., 2018); however, the role of *foxc1b* and its targets in the vasculature are not yet well understood.

Here, using a *foxc1b:EGFP* zebrafish transgenic reporter, we show that *foxc1b* labels smooth muscle cells and their precursors. We identify and visualize a key window for attachment of *foxc1b:EGFP* positive smooth muscle cells to the endothelium, and find smooth muscle differentiation is impaired in compound *foxc1a/foxc1b* mutants. The transcriptome of dual *foxc1b* + *acta2* expressing smooth muscle cells uncovers a set of enriched genes specific to the embryonic vascular smooth muscle cell population during early differentiation.

## Results

### *acta2, foxc1b*, and *pdgfrβ* label distinct and overlapping subsets of vascular mural cells in the developing embryo

Reliable markers to trace the development of the first vascular mural cells colonizing vessels are lacking. We therefore tested markers that are expressed in neural crest for whether they were also expressed in vascular smooth muscle around the ventral aorta, the first site to develop smooth muscle in the embryo. Such a marker would span the development of smooth muscle from when neural crest cells first attach to endothelium, to differentiation. *foxc1b*, as detected by the *foxc1b:EGFP* transgenic line, was one candidate. We first verified the transgenic line faithfully mimics endogenous *foxc1b* expression in mural cells. Comparing *foxc1b* mRNA expression at 2 and 4 dpf, patterns between the endogenous gene and transgenic line are very similar (Fig. 1A, C, E, G, I, and K). Specifically, *foxc1b* is expressed in the region of the aortic arches, jaw, ceratohyal (Schilling *et al*., 1996; Knight, 2003; Filipek-Górniok *et al*., 2015; Xu *et al*., 2018) and in the periocular mesenchyme in the ventral head. Furthermore, *foxc1b* mRNA expression overlaps with *foxc1b:EGFP* transgenic expression in the ventral aorta, aortic arch arteries, and ceratohyal at 2 and 4 dpf, suggesting EGFP in the transgenic fish line mimics endogenous mRNA expression (Fig. 1B, D, F, H, J, and L).

**Figure 1:**
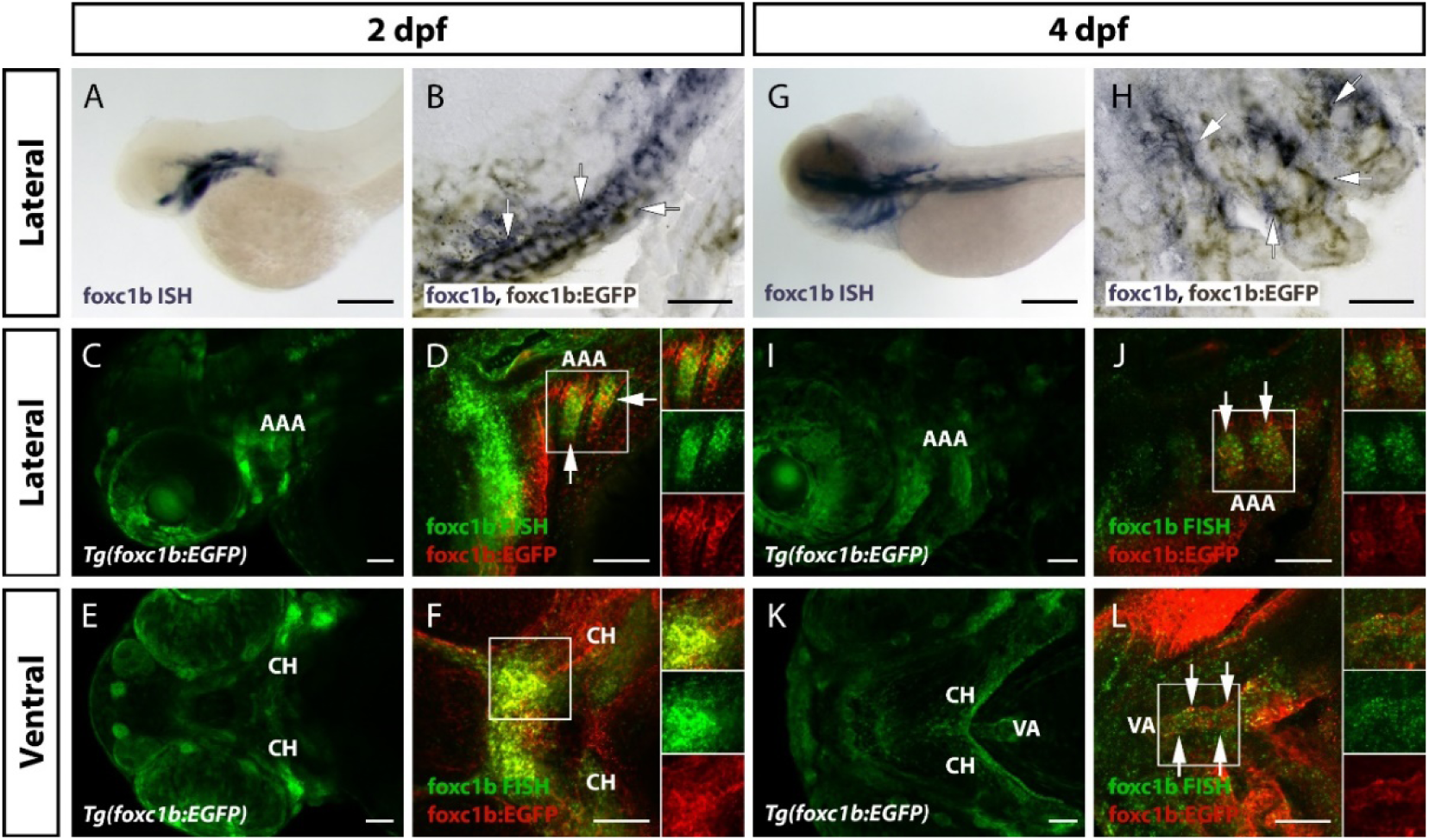
A *foxc1b:EGFP* transgenic line mimics endogenous *foxc1b* mRNA expression in the zebrafish ventral head. A-F) 2 dpf and G-L) 4 dpf images of *foxc1b* expression in lateral (A-D, G-J) and ventral views (E, F, K, L). A) *foxc1b* mRNA (purple) is expressed in the aortic arches, jaw, ceratohyal, and periocular mesenchyme of the ventral head of the zebrafish embryo. B) *foxc1b* mRNA and *foxc1b:EGFP* transgene expression (brown) overlap (arrows) in the ceratohyal. C) *foxc1b:EGFP* transgene (green) is strongly expressed in the aortic arches and mesenchyme of the ventral head. D) *foxc1b* mRNA expression (green) overlaps with *foxc1b:EGFP* antibody stain (red) in the aortic arch arteries (arrows). E) Strong expression of *foxc1b:EGFP* in the ventral head including the ceratohyal. F) *foxc1b* mRNA expression overlaps with *foxc1b:EGFP* antibody stain in the ceratohyal. G) At 4 dpf, *foxc1b* mRNA is present in the aortic arch arteries, the epibranchial region of the ventral brain, ceratohyal, swim bladder, and intestine. H) *foxc1b* mRNA and *foxc1b:EGFP* overlap in cells surrounding the aortic arch arteries. I) *foxc1b:EGFP* is expressed in aortic arch arteries. J) *foxc1b* mRNA expression overlaps with *foxc1b:EGFP* antibody stain in the aortic arch arteries (arrows). K) *foxc1b:EGFP* is expressed around the ventral aorta and ceratohyal. L) *foxc1b* mRNA expression overlaps with *foxc1b:EGFP* antibody stain in the ventral aorta (arrows) and ceratohyal. Scale bars in A and G represent 200 µm. Scale bars in B and H represent 20 µm. Scale bars in C-F and I-L represent 50 µm. Insets in D, F, J, and L show merged, *foxc1b:EGFP* F-ISH, and *foxc1b:EGFP* antibody stain. AAA = aortic arch arteries, CH = ceratohyal, VA = ventral aorta.

As Foxc1 is expressed in mural cells of mice (Siegenthaler *et al*., 2013), and *foxc1b* is expressed in endothelial cells of the early zebrafish trunk (Skarie and Link, 2009; Chen *et al*., 2017), we next tested the cell-type specificity of *foxc1b:EGFP* expression later, at 4 dpf, when robust expression of smooth muscle and pericyte markers are present in zebrafish embryos (Santoro, Pesce and Stainier, 2009; Seiler, Abrams and Pack, 2010; Wang *et al*., 2014; Whitesell *et al*., 2014). *foxc1b:EGFP* positive cells are located adjacent and in close contact with the endothelium of the ventral aorta (Fig. 2A, arrowheads) similar to cells expressing the classic smooth muscle marker, *acta2:EGFP* (Fig. 2B). We show that *foxc1b:EGFP* and *acta2:mCherry* are co-expressed in smooth muscle cells (Fig. 2C, arrows, arrowheads point to mural cells expressing only one marker). Similarly, when we compare *foxc1b* mRNA expression to *kdrl:EGFP* or *acta2:EGFP* transgenic fish we see *foxc1b* mRNA expressing cells are close to the endothelium (*kdrl*) and overlap with smooth muscle (*acta2*), but not the endothelium (Fig. S1). To test if *foxc1b:EGFP* also labels pericytes, *foxc1b:EGFP* was crossed with a *pdgfrβ:Gal4FF; UAS:NTR-mCherry* line (hereafter *pdgfrβ:mCherry*). At 4 dpf, we observed no co-expression of *foxc1b:EGFP* and *pdgfrβ:mCherry* in the ventral head (Fig. 2D). *pdgfrβ:mCherry* expressing cells are pericytes and are located adjacent to *kdrl:EGFP* expressing endothelial cells, and only rarely co-express *acta2:EGFP* (Fig. 2E, F).

**Figure 2:**
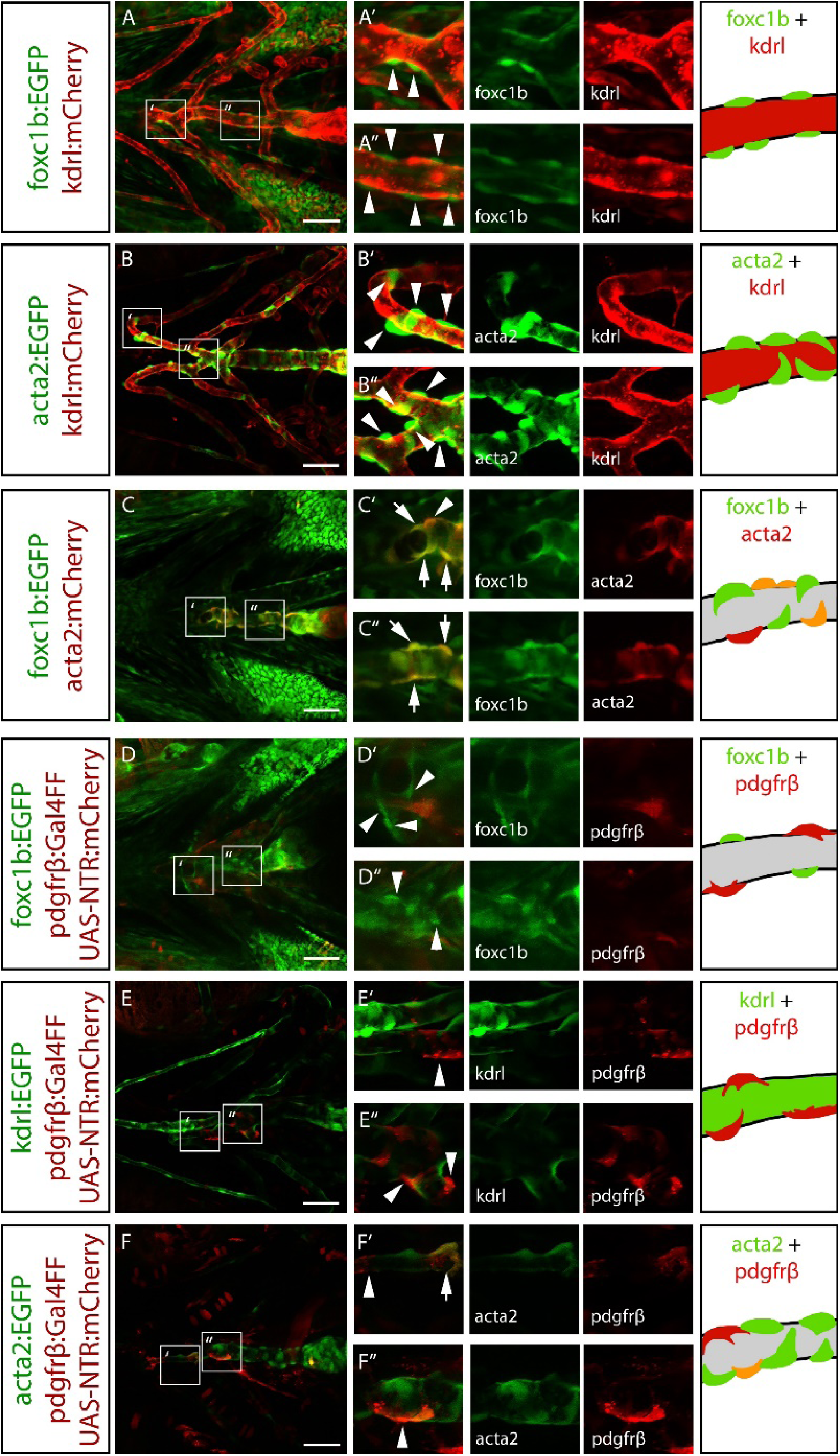
*foxc1b:EGFP* positive cells co-express a smooth muscle marker, but not endothelial or pericyte markers. Vessels of the ventral head in 4 dpf zebrafish. Arrowheads represent marker-expressing mural cells. Arrows show co-expression of two mural cell markers. A) *foxc1b:EGFP* expressing cells are associated with and surround the endothelium (*kdrl:mCherry*) along the ventral aorta. B) *acta2:EGFP* positive smooth muscle cells surround the *kdrl:mCherry* positive endothelium along the ventral aorta and aortic arch arteries. C) *foxc1b:EGFP* and *acta2:mCherry* are co-expressed along the ventral aorta. D) *foxc1b:EGFP* is not co-expressed in cells expressing a pericyte marker (*pdgfrβ:mCherry*) in the ventral head. E) *pdgfrβ:mCherry* is associated with the endothelium (*kdrl:EGFP*) in the ventral head. F) *pdgfrβ:mCherry* and *acta2:EGFP* are partially co-expressed in ventral head mural cells. Schematics depict mural cell and endothelial marker expression that match the transgenes. Grey indicates presumptive endothelial patterns. Scale bars represent 50 µm.

### *foxc1b* is expressed in mesenchymal cells as they associate with, and attach to, the endothelium

The earliest stages of migration and attachment of mural cells to the endothelial wall have not been observed in any species. We hypothesized that the early expression of *foxc1b* in neural crest and smooth muscle would allow us to visualize this process. At 2 dpf, when we have previously seen vascular mural cells present near the endothelium of the dorsal aorta using transmission electron microscopy (Liu *et al*., 2007; Lamont *et al*., 2010), *foxc1b:EGFP* positive cells are near the endothelium and have a mesenchymal morphology (Fig. 3A). In comparison, the *acta2:mCherry* transgene is not expressed in this region at this time (Fig. 3B) (Whitesell *et al*., 2014). By 3 dpf, *foxc1b:EGFP* positive cells associate closely with the endothelium (Fig. 3C, arrowheads) and some cells co-express *acta2:mCherry* (Fig. 3D, arrows). There are also *foxc1b:EGFP* positive cells that do not express *acta2:mCherry*. At 4 dpf, most mural cells co-express *foxc1b:EGFP* and *acta2:mCherry*. We note that when vascular expression of *acta2:mCherry* begins around 3 dpf in a few scattered cells, it is co-expressed with *foxc1b:EGFP* (Fig. 3D). Taken together, this suggests that *foxc1b* expression precedes *acta2* expression in mural cells (Fig. 3C, D). *foxc1b* also labels smooth muscle cells at later stages. At 4 dpf, *foxc1b:EGFP* is still present in mural cells around the endothelium (Fig. 3E), and these cells co-express *acta2:mCherry* (Fig. 3F). These patterns persist through 9 and 12 dpf (Fig. S2) and into adulthood. There is also co-expression of *foxc1b* and *acta2* in the trunk, along the dorsal aorta at 4 dpf, but no endothelial expression (Fig. S3). Co-expression is not universal, and cells expressing one or other marker (*foxc1b:EGFP* or *acta2:mCherry*) are occasionally observed (Fig. S2).

**Figure 3:**
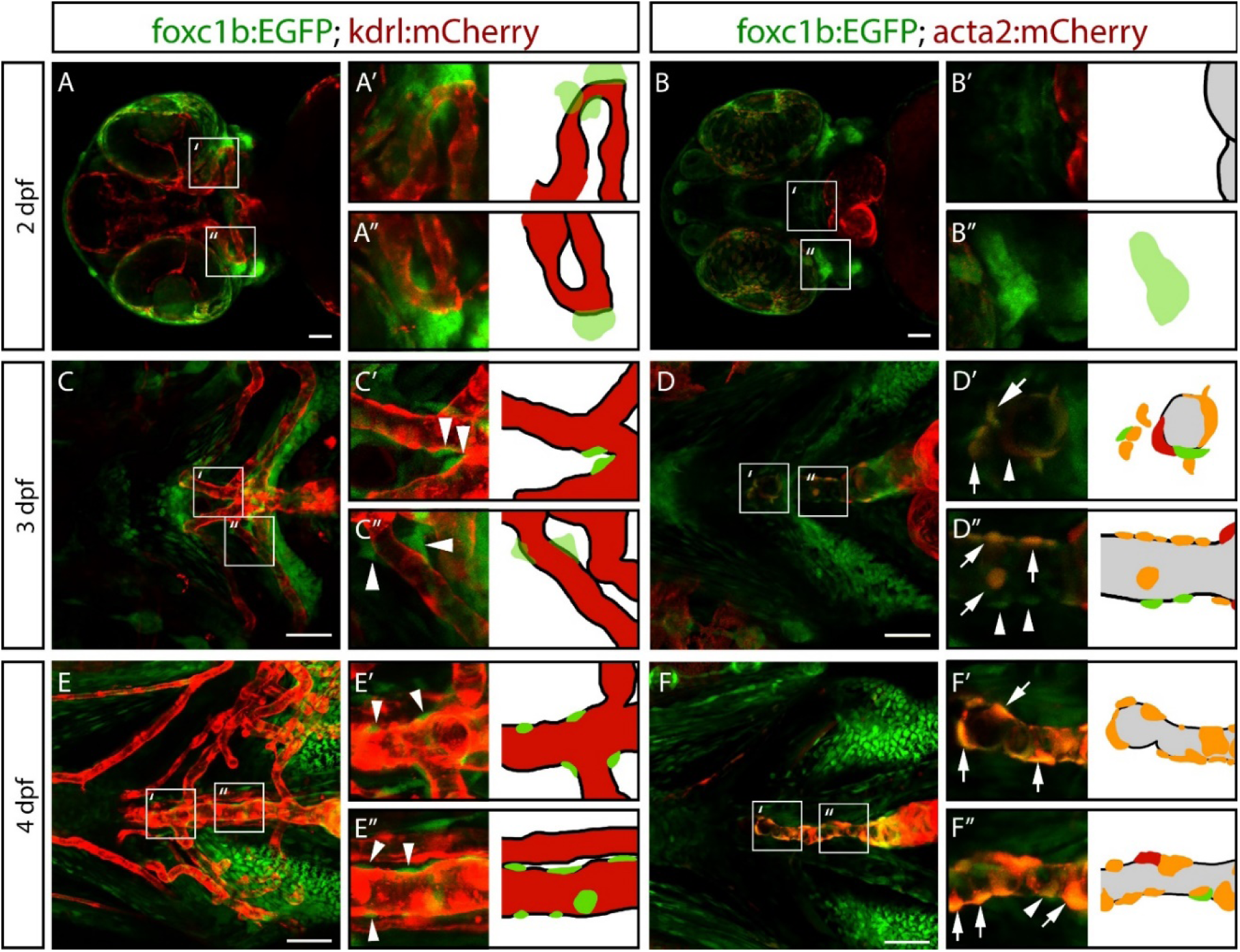
*foxc1b:EGFP* is expressed in mesenchymal cells as they associate with and attach to the endothelium. A, B) Images of the ventral head in 2 dpf zebrafish embryos show *foxc1b:EGFP* positive mesenchymal cells near the endothelium (*kdrl:mCherry*; n = 19 embryos), but not co-expressing a smooth muscle marker (*acta2:mCherry*; n = 6 embryos), which is only expressed in the heart at this stage. C, D) At 3 dpf, *foxc1b:EGFP* positive cells associate with the endothelium (n = 14 embryos), and some cells co-express *acta2:mCherry* (n = 5 embryos). Arrowheads indicate a mural cell marker associated with the endothelium. Arrows indicate co-expression between two mural cell markers. E, F) At 4 dpf, *foxc1b:EGFP* positive cells associate with the endothelium (n = 11 embryos) and a greater proportion of *foxc1b*:*EGFP* positive cells co-express *acta2:mCherry* (n = 8 embryos). Grey colour in schematics indicates endothelial patterns. Scale bars represent 50 µm.

To trace when *foxc1b:EGFP* expressing cells transform from a mesenchymal to a smooth muscle morphology we used time-lapse confocal microscopy. *foxc1b:EGFP* positive cells begin as mesenchymal clusters (Fig. 4A), adjacent to *kdrl:mCherry* positive endothelium. Distinct *foxc1b:EGFP* positive vascular mural cells emerge on the endothelium by 58 hpf (Fig. 4A, Movie 1). After taking on a mural cell morphology, *foxc1b:EGFP* positive cells then express *acta2*. This transition from *foxc1b:EGFP+; acta2:mCherry* to *foxc1b:EGFP+; acta2:mCherry+* cells occurs around 66 hpf (Fig. 4B, Movie 2). Taking time-lapse and marker expression together, our data depict the developmental progression for mural cells around the ventral aorta, where *foxc1b* expressing mesenchymal cells closely associate with the endothelium, morphologically differentiate into vascular mural cells and express the smooth muscle marker *acta2*.

**Figure 4:**
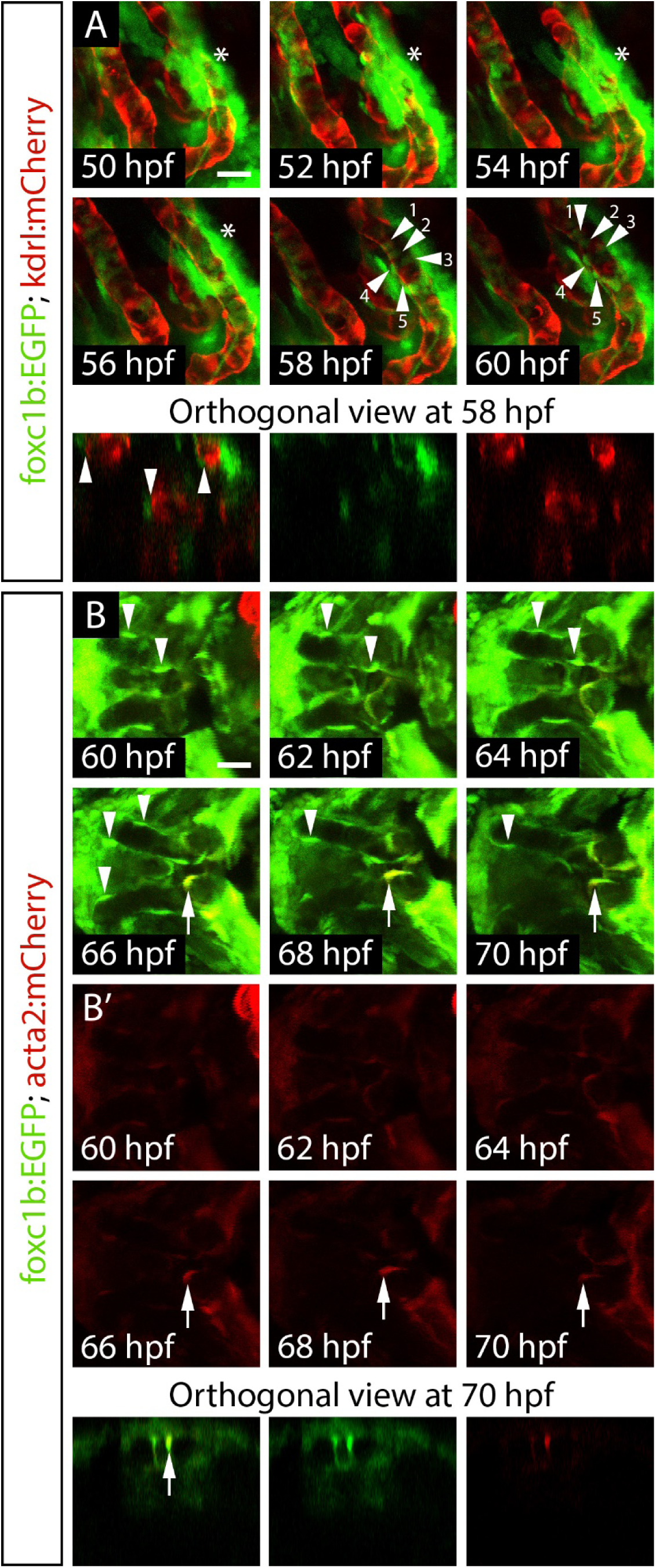
Time-lapse of *foxc1b:EGFP* positive mesenchymal cells associating with the endothelium and co-expressing a smooth muscle marker. A) Ventral views of aortic arch arteries of a *foxc1b:EGFP; kdrl:mCherry* embryo using confocal time-lapse imaging starting at 50 hpf. Mesenchymal cells are indicated by the asterisk, while vascular mural cells are indicated by arrowheads, starting by 56-58 hpf. Orthogonal views at 58 hpf depict the *foxc1b:EGFP* cells wrapping around, but not co-expressing an endothelial marker. B) Time-lapse of the early ventral aorta of a *foxc1b:EGFP; acta2:mCherry* embryo from 60-70 hpf. *foxc1b:EGFP* positive smooth muscle cells are associated with the endothelium (arrowheads) but do not co-express *acta2:mCherry* until approximately 66 hpf (arrow). B’) Single channel images of *acta2:mCherry.* Orthogonal views at 70 hpf show *acta2:mCherry* co-expressed within a *foxc1b:EGFP* positive cell. Scale bars represent 20 µm.

### *foxc1b* is expressed in smooth muscle cells, but not pericytes, in the adult zebrafish brain

Foxc1 marks both pericytes and smooth muscle cells in the mouse brain, but it is unknown whether it shows cell-type specificity of expression in zebrafish at adult stages. We find that embryonic co-expression patterns of *foxc1b, acta2* and *pdgfrβ* at 4 dpf are consistent in the adult brain, but also show some important differences. In the adult brain, we find that *foxc1b:EGFP* expressing cells are associated with large arterioles, but not smaller capillaries (Fig. 5A), similar to *acta2:EGFP* expressing cells (Fig. 5B). *foxc1b:EGFP* is almost always co-expressed with *acta2:mCherry*; however, there are regions of *acta2:mCherry* expression that lack co-expression of *foxc1b:EGFP* (Fig. 5C), particularly on the largest diameter vessels. We never detected *foxc1b:EGFP* expressing cells on the same vessels that express *pdgfrβ:mCherry*, and *foxc1b:EGFP* is not expressed in endothelial cells (Fig. 5A). Thus, in both the ventral head of the embryo and the adult brain, *foxc1b* is expressed in smooth muscle cells, but not pericytes or endothelial cells.

**Figure 5:**
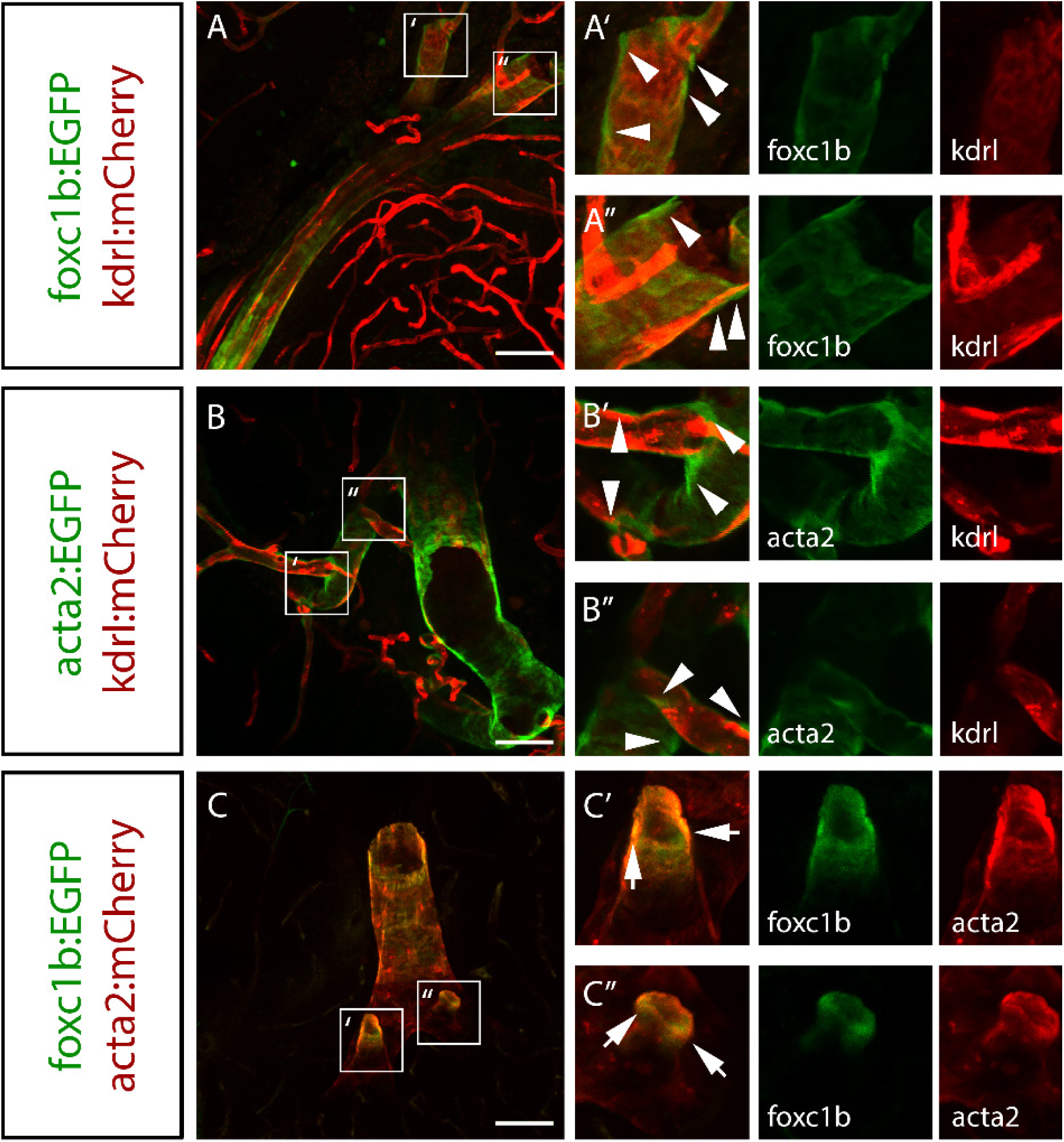
*foxc1b:EGFP* expression partially overlaps with *acta2* expression in smooth muscle cells in the adult zebrafish brain. A) *foxc1b:EGFP* expressing cells are closely associated with endothelial cells (*kdrl:mCherry*) in the adult brain. Arrowheads point to mural cells associated with the endothelium. Arrows point to co-expression of two mural cell markers. B) *acta2:EGFP* expressing smooth muscle cells are closely associated with *kdrl:mCherry* expressing endothelial cells along large vessels of the brain. C) *foxc1b:EGFP* and *acta2:mCherry* are partially co-expressed in smooth muscle cells on brain vessels. Scale bars represent 50 µm.

### Mural cell marker expression changes with vessel diameter and location

As an animal grows, its vessels enlarge to increase the supply of blood to growing tissues. We expected that mural cell types would correlate with vessel diameter, but that there might also be some variation in mural cell type depending on vessel location and developmental timing. We measured vessel diameter in vessels expressing *kdrl* (no mural cells), *pdgfrβ, foxc1b* or *acta2* in the ventral head (ventral aorta, opercular artery, hypobranchial artery, aortic arch arteries, and lateral dorsal aorta), embryonic brain (central arteries, primordial hindbrain channel, basilar artery, and the internal carotid artery), and adult brain.

Starting at 4dpf, when *acta2* is first expressed in smooth muscle cells of the ventral aorta and associated vessels, we find that naked vessels (labelled by endothelial *kdrl*, but unlabelled with mural cell markers) have a mean diameter of 9.7 ± 2.0 µm (Fig. 6A, B). The vessels covered with pericytes (*pdgfrβ*) have a mean vessel diameter of 10.3 ± 2.1, but are not significantly different in size from naked vessels. We find the mean vessel diameter for *foxc1b* covered vessels is 14.6 ± 4.4 µm, while for *acta2* covered vessels, the mean is 14.1 ± 4.5 µm, and for *foxc1b + acta2* covered vessels the mean diameter is 20.5 ± 6.8 µm. Thus, vessels with *foxc1b* or *acta2* single positive mural cells are significantly larger than naked *kdrl* or *pdgfrβ* covered capillaries, as one might expect for vessels covered by smooth muscle, and *foxc1b* + *acta2* double marked vessels are significantly larger. The reason for larger double positive vessels is unclear, but may reflect a strong contribution of *foxc1b* positive cells to the ventral aorta where *acta2* is heavily expressed in early development.

**Figure 6:**
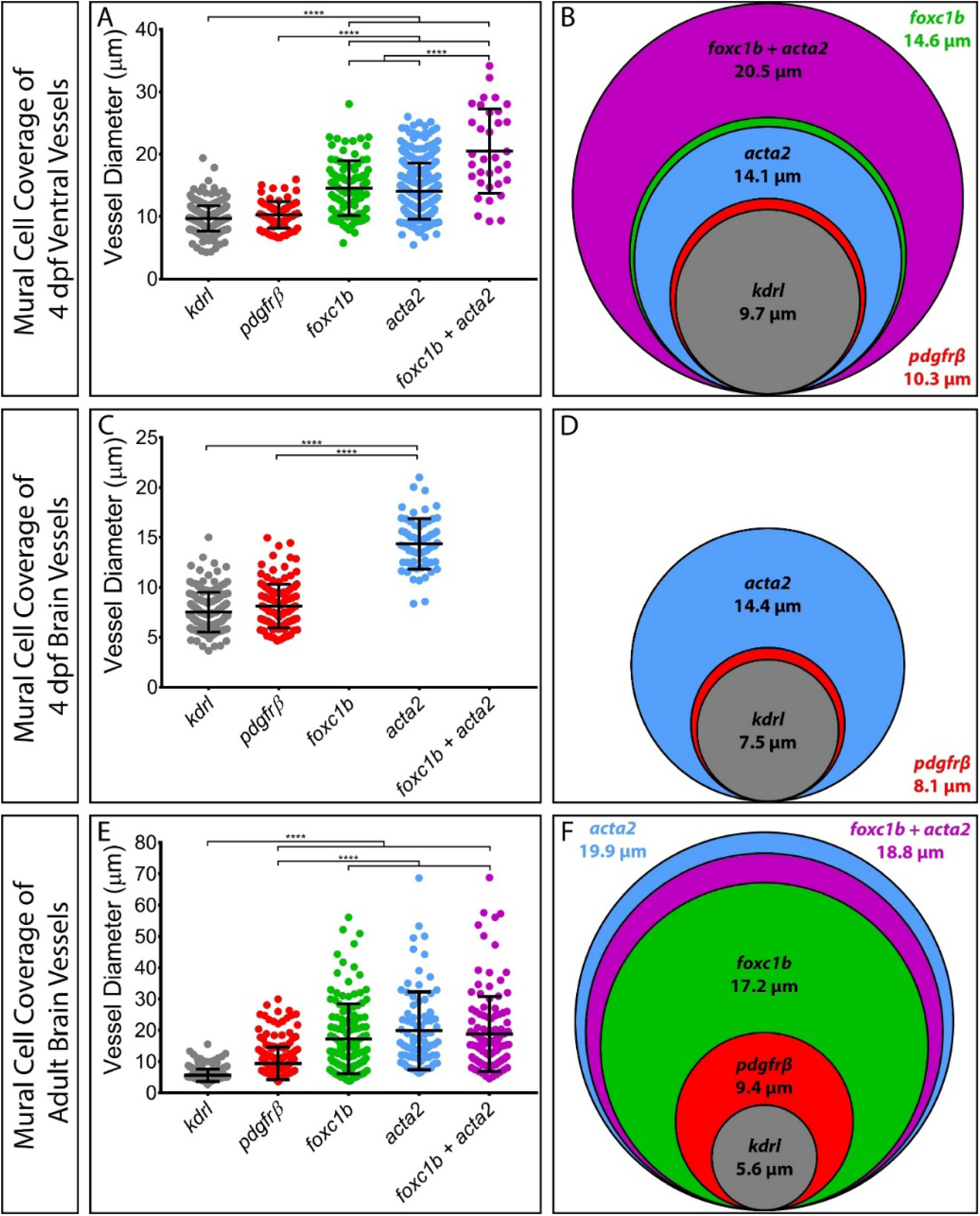
Mural cell marker expression is dependent upon vessel diameter, location, and developmental stage. A) Vessel diameters were measured from the endothelium adjacent to mural cell markers at in the ventral head. Naked vessels are vessels labelled by the endothelial *kdrl* transgene only, while vessels covered in mural cells are categorized by the combination of mural cell markers expressed. Measurements of 4 dpf ventral head vessels: *kdrl* (9.7 ± 2.0 µm, n=357), *pdgfrβ* (10.3 ± 2.1 µm, n=75), *foxc1b* (14.6 ± 4.4 µm, n=89), *acta2* (14.1 ± 4.5 µm, n=308), and *foxc1b* + *acta2* (20.5 ± 6.8 µm, n=34). B) Schematic of mean vessel diameter per condition with vessel sizes scaled relative to each other. C) Measurements of 4 dpf embryonic brain vessels: *kdrl* (7.5 ± 2.0 µm, n=147), *pdgfrβ* (8.1 ± 2.2 µm, n=119), and *acta2* (14.4 ± 2.5 µm, n=63). D) Schematic of mean vessel diameter per condition with vessel sizes scaled relative to each other. E) Measurements of adult brain vessels: *kdrl* (5.6 ± 2.0 µm, n=244), *pdgfrβ* (9.4 ± 5.2 µm, n=238), *foxc1b* (17.2 ± 11.2 µm, n=133), *acta2* (19.9 ± 12.5 µm, n=83), and *foxc1b* + *acta2* (18.8 ± 12.0 µm, n=126). F) Schematic of mean vessel diameter per condition with vessel sizes scaled relative to each other. **** = p < 0.0001, ANOVA with Tukey’s multiple comparisons test.

**Figure 7:**
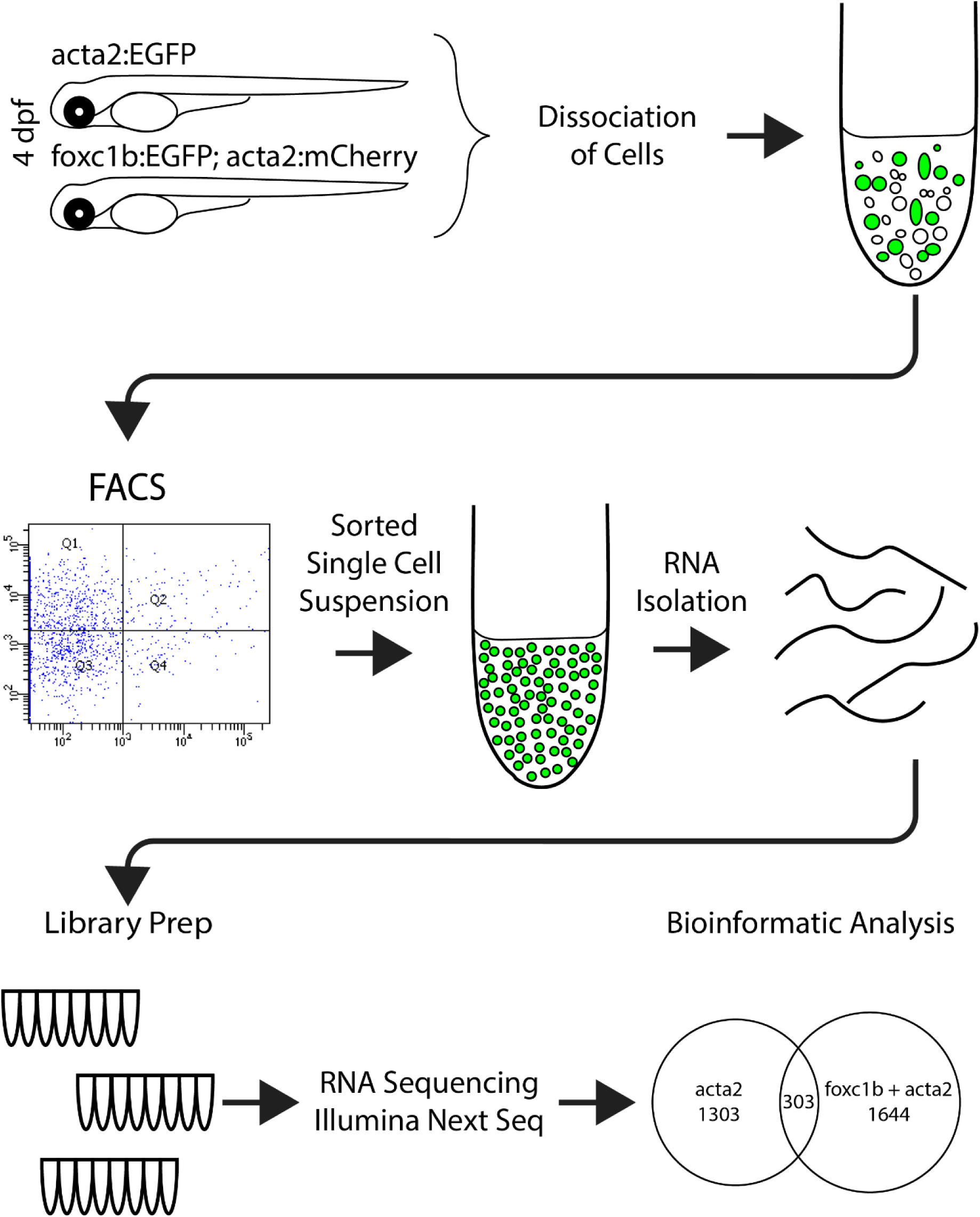
Workflow for isolating smooth muscle transcriptomes from zebrafish embryos. 4 and 5 dpf zebrafish embryos were pooled and dissociated before fluorescence-activated cell sorting. RNA was isolated and sequenced using an Illumina Next Seq platform.

The embryonic brain develops smooth muscle coverage later than vessels in the ventral head, most likely as the blood flow in the brain is lower in early development. Interestingly, we found no expression of *foxc1b* around vessels in the 4 dpf zebrafish brain. The mean vessel diameter for naked (*kdrl*) brain vessels at 4 dpf is 7.5 ± 2.0 µm while it is 8.1 ± 2.2 µm for *pdgfrβ* covered vessels (Fig. 6C, D). Both naked *kdrl* and *pdgfrβ* covered vessel diameters are significantly smaller than *acta2* covered vessels (14.4 ± 2.5 µm).

In the adult brain, the mean vessel diameter for naked vessels is 5.7 ± 2.0 µm and for *pdgfrβ* covered vessels it is significantly larger at 9.4 ± 5.2 µm. The mean vessel diameter of *foxc1b* covered vessels is 17.2 ± 11.2 µm, and for *acta2* covered vessels the diameter is 19.9 ± 12.5 µm. For vessels that express both *foxc1b* + *acta2,* the diameter is 18.8 ± 12.0 µm (Fig. 6E, F). Overall, vessels covered by *foxc1b* are significantly larger than both naked and *pdgfrβ* covered vessels, but all smooth muscle covered vessels (*foxc1b, acta2, foxc1b + acta2*) are of a similar size. Representative vessel sizes are depicted, to scale in Figure 6F. Thus we show that *foxc1b:EGFP* labels smooth muscle cells on medium and large diameter vessels in the adult brain.

### Mural cell populations have unique transcriptome profiles

To probe the transcriptome of vascular smooth muscle cells in early development, we required a specific marker; however, expression of the *acta2:EGFP* transgene is not restricted to vascular smooth muscle cells, limiting its utility for transcriptome profiling. Thus, the intersection of *foxc1b* and *acta2* transgene expression might provide the possibility for more selective smooth muscle cell isolation. To investigate this possibility, we isolated *acta2:EGFP* positive or *foxc1b:EGFP + acta2:mCherry* double positive cells, along with respective negative populations, and subjected them to RNA-Seq. In parallel, we also analyzed *kdrl:HRAS-mCherry* positive and negative cells for comparison. Consistent with a mural cell identity, both *acta2:EGFP* positive and *foxc1b:EGFP + acta2:mCherry* double positive cells show significant enriched expression of known vascular smooth muscle genes, such as *acta2, tagln* and *fbln5*, when compared to transgene negative populations (Fig. 8A, B). By contrast, neither population showed an enrichment for endothelial markers (*cdh5, pecam1, egfl7*; Fig. 8A-C). Interestingly, *myocardin* (*myoc*), a known transcriptional regulator of smooth muscle cell differentiation in mammals (Du *et al*., 2003) only showed significant enrichment when comparing *foxc1b:EGFP + acta2:mCherry* double positive cells versus negative (Fig. 8A, B), suggesting that intersectional transgene labeling improved detection of known mural cell genes. Indeed, multiple vascular smooth muscle cell expressed genes showed much higher levels of enrichment in *foxc1b:EGFP + acta2:mCherry* double positive cells when compared to negative cells, than *acta2:egfp*-positive (Fig. 8D). As noted above, all but one of these genes failed to show any enrichment in endothelial cells (Fig. 8D). To further investigate the utility of intersectional labeling of mural cell populations using *foxc1b:EGFP* and *acta2:mCherry*, we performed whole mount *in situ* hybridization to validate candidate genes’ expression patterns. For this purpose, we chose 11 candidate transcripts whose expression was enriched in *foxc1b:EGFP + acta2:mCherry* double positive cells (Fig. 8E, Fig. 9). In all cases, we observed expression in putative mural cells surrounding the endothelial lining of blood vessels. Moreover, two candidate genes (*capn3b* and *cracr2b/si:ch211-247i17.1*) only displayed enrichment in *foxc1b:EGFP + acta2:mCherry* double positive cells, and exhibit expression in putative mural cells. Together, these observations demonstrate the potential utility for using an intersectional approach to label vascular mural cell populations. Moreover, this approach aids in the identification of potentially new candidate genes that may play a role in mural cell development or function.

**Figure 8:**
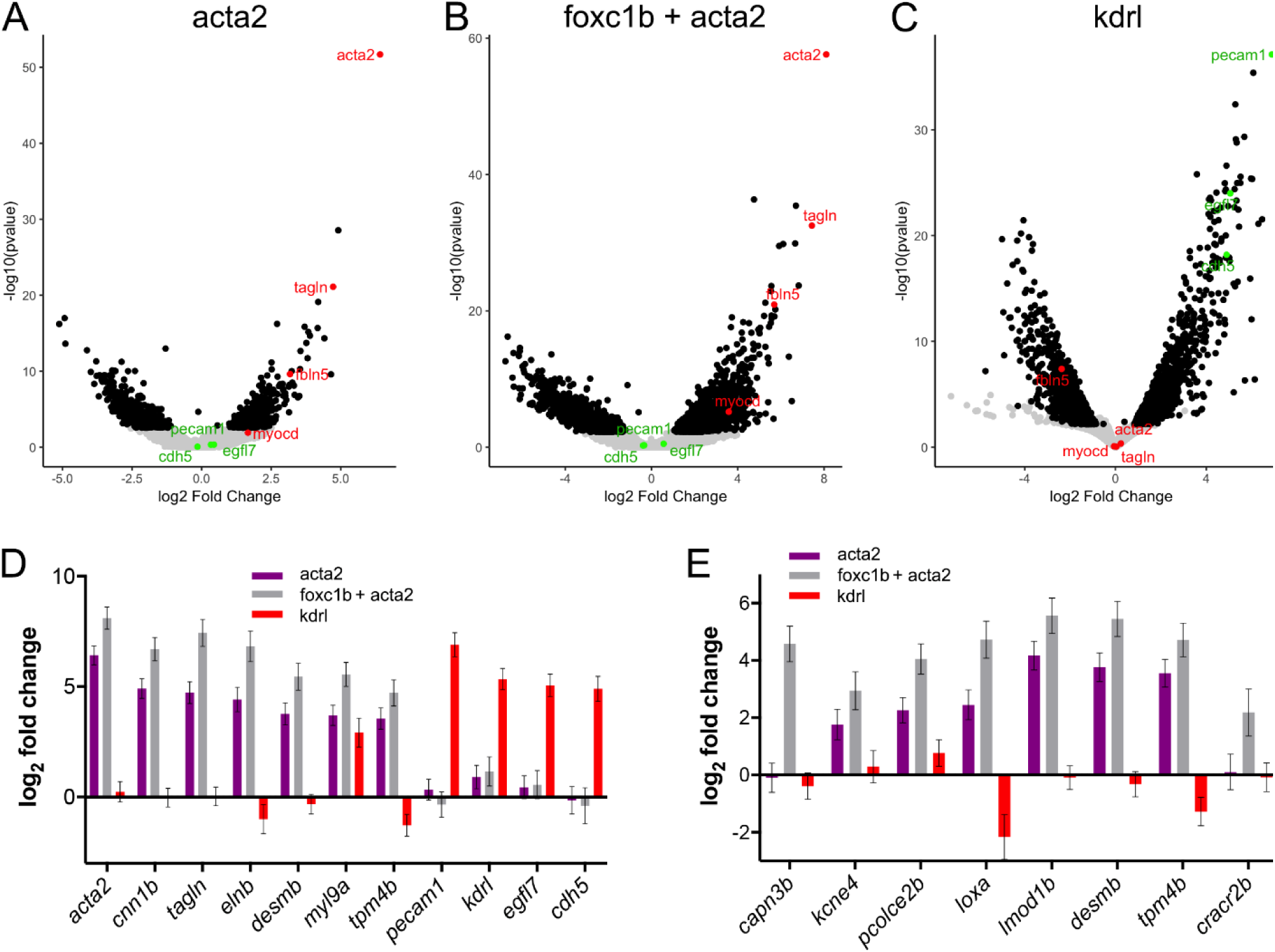
Differential gene expression in embryonic smooth muscle cells. A-C) Volcano plots for *acta2* smooth muscle (A), *foxc1b + acta2* smooth muscle (B), and endothelial (C) datasets. Representative smooth muscle genes are shown in red, whereas endothelial genes are shown in green. Black dots represent genes with significant changes (p adj < 0.05). D) Log_2_ fold change in each dataset for genes indicated in the volcano plots. E) Log_2_ fold change in each dataset for some genes indicated in Supplemental Figure 5. Error bars in D and E are standard error.

**Figure 9:**
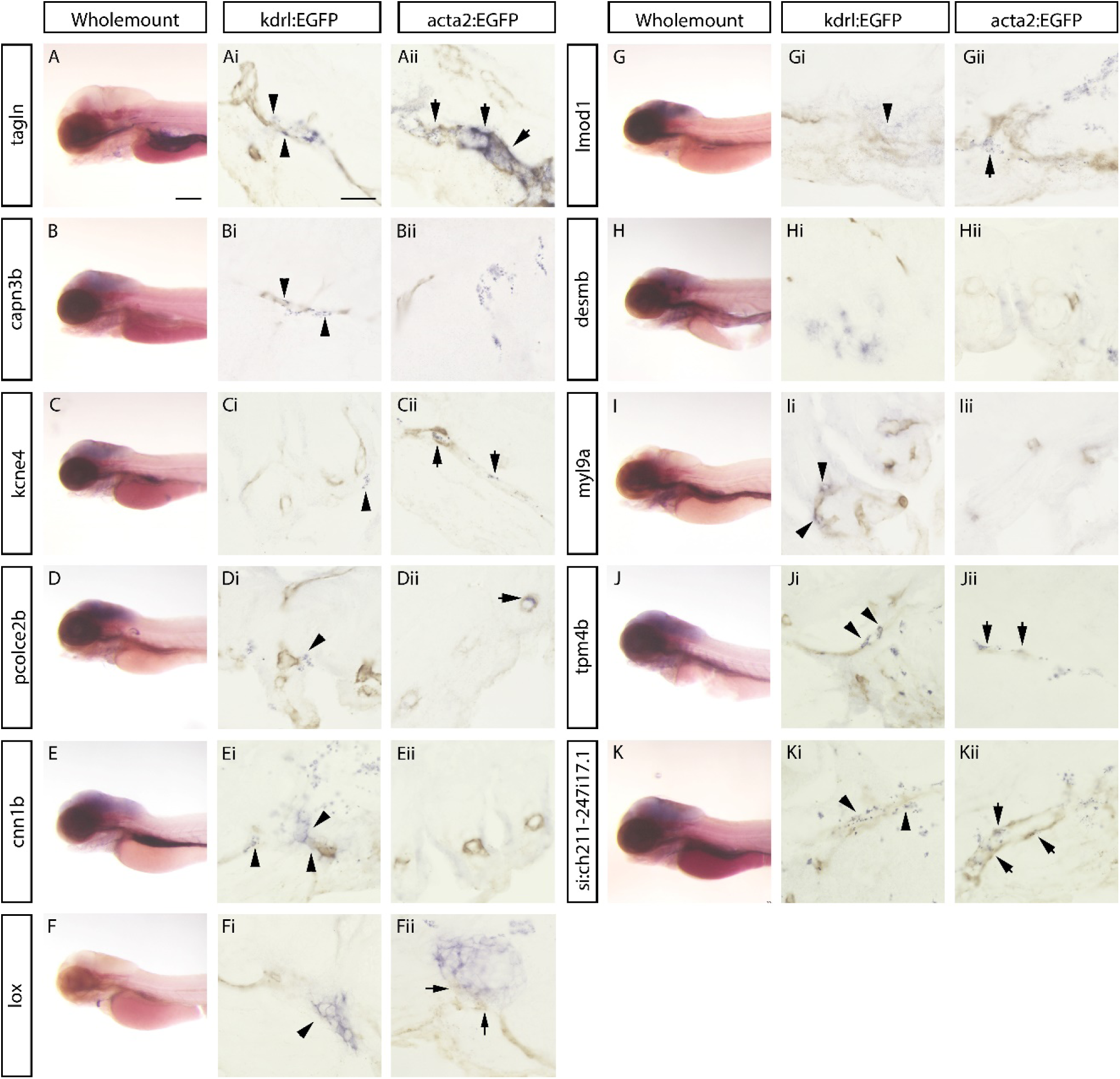
Validation of genes from the mural cell transcriptomes. Gene expression in lateral view wholemount and sections, compared to endothelial (antibody staining for *kdrl:EGFP*, brown, column i), or smooth muscle (*acta2:EGFP*, brown, column ii) transgenes. Arrowheads depict gene expression associated with the endothelium, arrows depict gene expression co-expressed with smooth muscle. A) *tagln*, a known smooth muscle marker, is found in the aortic arch arteries, notochord tip, swim bladder, and gut in wholemount. In sections *tagln* is expressed in cells adjacent to the endothelium and is co-expressed with *acta2:EGFP* on the ventral aorta and aortic arch arteries. B) *capn3b* is expressed in cells adjacent to the endothelium in the brain. C) *kcne4* is expressed adjacent to the endothelium and is co-expressed in smooth muscle on the ventral aorta and aortic arch arteries. D) *pcolce2b* is expressed adjacent to the endothelium and is co-expressed with *acta2:EGFP*. E) *cnn1b* is expressed adjacent to the endothelium of the aortic arch arteries. F) *lox* is expressed solely in the smooth muscle of the bulbus arteriosus. G) *lmod1* is expressed adjacent to the endothelium and overlaps with *acta2:EGFP*. H) *desmb* is expressed in a punctate pattern in the brain. I) *myl9a* is expressed adjacent to the endothelium, and partially overlaps with *acta2:EGFP*. J) *tpm4b* is expressed near the endothelium in the ventral head and is co-expressed with *acta2:EGFP* in the brain. K) *si:ch211-247i17.1* is expressed near the endothelium and is co-expressed with *acta2:EGFP*. Scale bar for A-K (wholemount) represent 200 µm. Scale bars for sections (i and ii) represent 20 µm.

### *foxc1* genes are required for smooth muscle development

As *foxc1b* is expressed in cells that differentiate into *acta2* positive smooth muscle cells, we next tested whether *foxc1b* and its paralog *foxc1a* are necessary for smooth muscle development. The two genes have similar expression patterns, and *foxc1a* and *foxc1b* can compensate for each other (Jolanta M. Topczewska *et al*., 2001; Skarie and Link, 2009; Xu *et al*., 2018). *foxc1a* is highly expressed in our *acta2* dataset, thus we scored for *acta2:EGFP* positive smooth muscle cell coverage of the ventral aorta in *foxc1a* and *foxc1b* compound mutants. Incrosses of *foxc1a*^*+/-*^; *foxc1b*^*+/-*^ compound heterozygote zebrafish provided all 9 genotype combinations (Fig. 10A-I). In addition to the mutant background, all embryos were heterozygous for the *acta2:EGFP* transgene to visualize smooth muscle. Wild type *foxc1a*^*+/+*^; *foxc1b*^*+/+*^ embryos have a mean length of smooth muscle coverage of (102.3 µm). Loss of *foxc1b* alleles in a *foxc1a*^*+/+*^ background causes no significant change in coverage (*foxc1a*^*+/+*^; *foxc1b*^*+/-*^ = 134.7 µm, *foxc1a*^*+/+*^; *foxc1b*^*-/-*^ = 94.0 µm, Fig. 10A – C). There is also no significant change with heterozygous loss of *foxc1a*^*+/-*^ (*foxc1a*^*+/-*^; *foxc1b*^*+/+*^ = 99.5 µm, *foxc1a*^*+/-*^; *foxc1b*^*+/-*^ = 101.9 µm, and *foxc1a*^*+/-*^; *foxc1b*^*-/-*^ = 83.4 µm; Fig. 10D – F). However, homozygous loss of *foxc1a*^*-/-*^ has a strong effect on smooth muscle coverage when combined with loss of *foxc1b*. *foxc1a*^*-/-*^; *foxc1b*^*+/+*^ embryos are not significantly different from wild type (94.1 µm). In contrast, *foxc1a*^*-/-*^; *foxc1b*^*+/-*^ (3.2 µm), and *foxc1a*^*-/-*^; *foxc1b*^*-/-*^ (13.1 µm; Fig. 10G– I) embryos have severely reduced smooth muscle coverage on the ventral aorta. Thus, the length of smooth muscle coverage along the ventral aorta decreases with increased loss of *foxc1* alleles, with loss of 3 or 4 alleles having the most significant reduction in the length of smooth muscle coverage (Fig. 10J).

**Figure 10:**
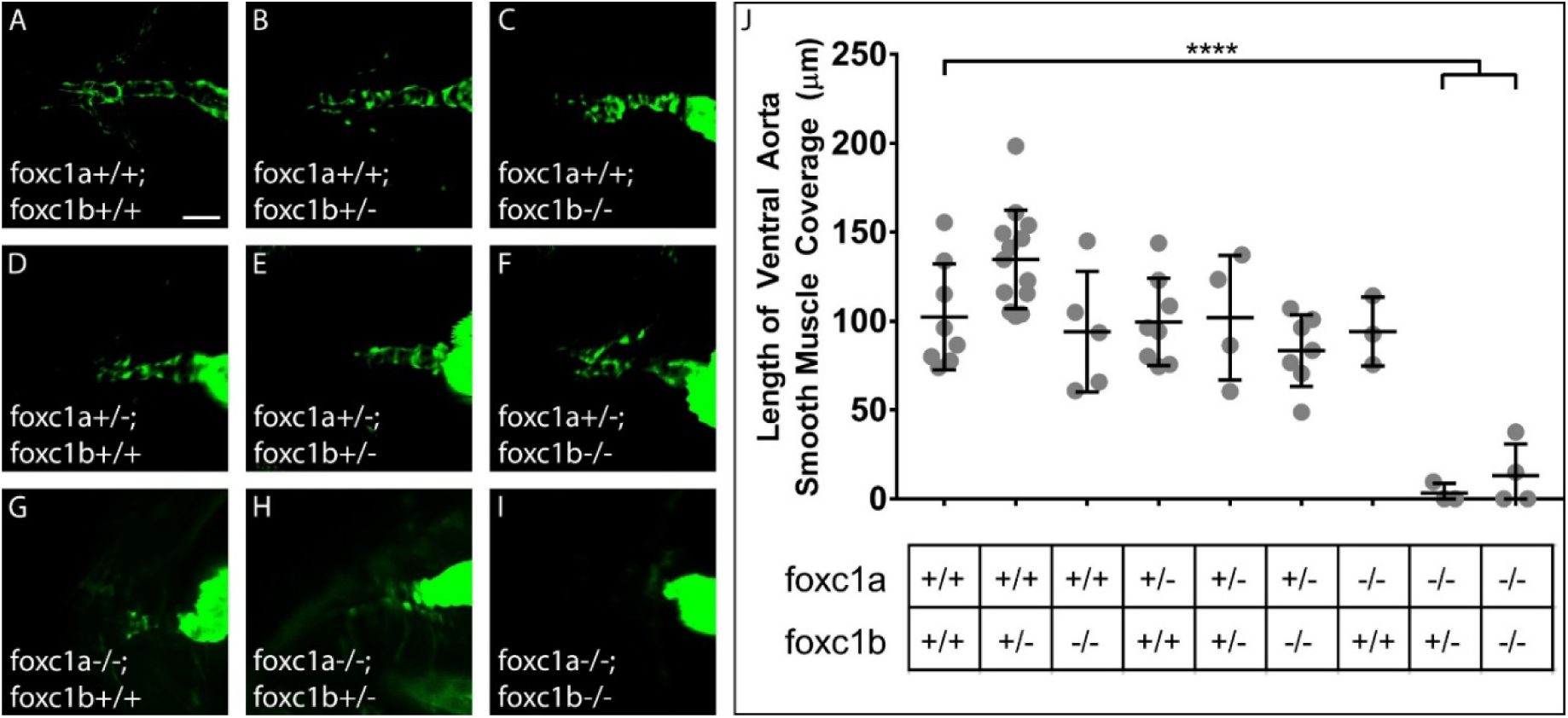
*foxc1a/foxc1b* mutants show increased loss of smooth muscle coverage with the progressive loss of *foxc1a/foxc1b* alleles. Smooth muscle coverage (*acta2:EGFP*) at 4 dpf on the ventral aorta of *foxc1a* and *foxc1b* compound mutants. Genotypes are listed in each corresponding panel. A-C) *foxc1a*^*+/+*^ fish show no statistical change in the length of ventral aorta smooth muscle coverage, with progressive loss of *foxc1b*^*(+/+ > +/-> -/-)*^. D-F) *foxc1a*^*+/-*^ mutants show no statistical change in the length of ventral aorta smooth muscle coverage with progressive loss of *foxc1b*^*(+/+ > +/-> -/-)*^. G-I) *foxc1a*^*-/-*^ mutants show a statistical significant decrease in *acta2:EGF*P positive cells on the ventral aorta in *foxc1b*^*+/-*^ and *foxc1b*^*-/-*^ mutants. J) Quantification of the length of ventral aorta smooth muscle coverage, **** = p < 0.0001, ANOVA with Dunnett’s multiple comparisons test. Scale bar represents 50 µm.

Of note, when both alleles of *foxc1a* are lost, these embryos initially develop circulation but present with progressive cardiac edema and circulatory insufficiency around 72 hpf (Fig. S5). These data suggest that not only is *foxc1* expressed early in the smooth muscle lineage, but that *foxc1 (a and b*) are critical for the early development of smooth muscle cells, and cardiac function. Homozygous loss of *foxc1a* coupled with loss of one or two alleles of *foxc1b* has the strongest effect on vascular development, but the converse is not true. Loss of two alleles of *foxc1b* and one allele of *foxc1a* has a wild type phenotype.

## Discussion

Vascular mural cells are required for maintenance, contraction, and support of the underlying endothelium; however, we still have limited tools to study their early development, *in vivo.* Current common markers of smooth muscle cells (*acta2, tagln)* and pericytes (*pdgfrβ, ng2*) have relatively specific and robust expression in the cells of interest but smooth muscle markers are not expressed early enough in development in the zebrafish model to study early mural cell differentiation. Furthermore, neural crest markers such as *sox10* turn off before smooth muscle cells differentiate in fish, unless utilizing lineage tracing techniques (Cavanaugh, Huang and Chen, 2015). However, we know that neural crest cells are precursors of vascular smooth muscle and pericytes in zebrafish (Cavanaugh, Huang and Chen, 2015; Ando *et al*., 2016; Stratman *et al*., 2017), and we can identify vascular mural cells using transmission electron microscopy before we observe expression of smooth muscle markers (Liu *et al*., 2007; Lamont *et al*., 2010). Identification of additional markers is therefore necessary to observe these intermediate differentiation states. We show here *foxc1b:EGFP* labels early mesenchymal cells and smooth muscle cells *in vivo* and we can observe these cells undergoing morphological changes from a mesenchymal to a smooth muscle morphology. We use *foxc1b* to study morphological differentiation of smooth muscle cells in real time, the interplay between mural cell marker expression, vessel size and its transcriptome, as well as the role of *foxc1* in smooth muscle differentiation. Furthermore, selecting a *foxc1b + acta2* positive subset of mural cells identifies an enriched vascular mural cell population.

### *foxc1b* is involved in early vascular smooth muscle cell differentiation

We first established that *foxc1b* expressing cells co-express a smooth muscle marker, *acta2,* as in mouse (Kume *et al*., 2001; Siegenthaler *et al*., 2013). Unlike mouse, *foxc1b:EGFP* is not co-expressed in head endothelial cells or pericytes, suggesting that some *foxc1b:EGFP* expressing cells are, or will become smooth muscle cells in zebrafish. Three lines of evidence suggest that *foxc1b* is expressed in mesenchymal precursors that will become *acta2* positive smooth muscle cells. First, we observe mesenchymal expression of *foxc1b:EGFP* near vessels at 2 dpf and show how these cells undergo morphogenesis as they adhere to endothelium and adopt a smooth muscle morphology. Secondly, *foxc1b* is expressed in vascular mural cells before the earliest *acta2* expression, and at later timepoints, the two genes are strongly co-expressed. Thirdly, loss of *foxc1* leads to a strong reduction in the number and the coverage by *acta2* positive cells on the ventral aorta. Taken together, these data suggest *foxc1b* expressing mesenchymal precursors differentiate into *acta2* expressing vascular smooth muscle cells on the ventral aorta.

In mice, FOXC proteins are regulated by Sonic hedgehog (Shh), a signaling pathway driving mural cell differentiation (Yamagishi *et al*., 2003; Lamont *et al*., 2010). Given that *foxc1b* is expressed in neural crest (Seo and Kume, 2006; Inman *et al*., 2013; Koo and Kume, 2013; Seo *et al*., 2017), that lineage tracing of *sox10*-expressing neural crest cells show they contribute to smooth muscle on the bulbus arteriosus and ventral aorta (Mongera *et al*., 2013; Cavanaugh, Huang and Chen, 2015), and FOXC proteins are expressed in mural cells in mouse and fish (Kume *et al*., 2001; Siegenthaler *et al*., 2013; French *et al*., 2014), our data are consistent with previous models of the origin of the first vascular smooth muscle cells in the head of the embryo (Etchevers *et al*., 2001; Calloni *et al*., 2007; Mongera *et al*., 2013; Wang *et al*., 2014; Cavanaugh, Huang and Chen, 2015).

The role of *foxc1* in smooth muscle differentiation in zebrafish is unique. For instance, compound Foxc1 and Foxc2 mutant mouse vessels maintain *acta2* positive smooth muscle coverage, although vessel morphology is altered, with only enlarged vessels present (Kume *et al*., 2001). However, in fish we show that the loss of at least 3 alleles of *foxc1a/b* prevents *acta2* positive smooth muscle from differentiating along the ventral aorta. Similar to the morpholino knockdown of *foxc1a* and *foxc1b* (French *et al*., 2014), loss of both paralogs is required for significant loss of smooth muscle coverage, with *foxc1a* having a stronger effect on smooth muscle differentiation than *foxc1b*. There is known genetic redundancy in zebrafish *foxc1* genes (Jolanta M. Topczewska *et al*., 2001; Xu *et al*., 2018). *foxc1a* has a similar expression pattern, to *foxc1b* (Jolanta M. Topczewska *et al*., 2001; Xu *et al*., 2018) and Fox family members are known to play redundant roles in mice (Kume *et al*., 2001; Seo and Kume, 2006).

### Localization of mural cell subsets and the transcriptomic differences among embryonic mural cells

*foxc1* is expressed in multiple cell types in both the brain and trunk of mice and zebrafish. In the trunk of the zebrafish, *foxc1b* mRNA is expressed within endothelial cells during very early angiogenesis (Skarie and Link, 2009; Chen *et al*., 2017); however, we do not see transgenic *foxc1b:EGPF* expression in head or trunk endothelial cells at the late developmental stages we examine (4 dpf through adult) (Kume *et al*., 1998; Prasitsak *et al*., 2015). Surprisingly, despite reported strong expression of Foxc1 in mouse brain pericytes, we do not see *foxc1b* expression in fish brain pericytes at any stage (Zarbalis *et al*., 2007; Siegenthaler *et al*., 2013; Mishra *et al*., 2016). Instead we find that *foxc1b:EGFP* expression is within smooth muscle cells, with a similar expression pattern to *acta2.* Furthermore, we show that *foxc1a* and *b* expression is required at an early stage in the differentiation of *acta2* positive smooth muscle cells around the ventral aorta. These differences in expression may reflect species or developmental staging differences between fish and mammals, or incomplete elements in the enhancer used to generate the *foxc1b* transgene.

We wanted to understand whether *foxc1b* and *acta2* are always co-expressed and what type of vessels have mural cells expressing these markers. We chose to examine this by vessel diameter, a measurement that is relatively constant for capillaries across species. The smallest capillary vessel diameters are set by the size of the erythrocyte (about 7-8 μm in diameter in humans and zebrafish) (Kulkeaw and Sugiyama, 2012) and are therefore comparable cross-species (Hartmann *et al*., 2015). We found a complete lack of co-expression of *foxc1b* with *pdgfrβ*, a pericyte marker and therefore absence of *foxc1b* expressing cells from the smallest vessels of the vascular system. Instead, we found a hierarchical continuum of mural cell marker expression, with smooth muscle marked by *foxc1b* and *acta2* on larger diameter vessels. As the ventral aorta is the largest vessel in the embryonic ventral head, it is subject to larger blood pressure changes and needs the largest amount of smooth muscle coverage in the early developing embryo (Hu, Joseph Yost and Clark, 2001; Whitesell *et al*., 2014). Therefore, the need for more mature, contractile smooth muscle early in development as compared to the brain, may require early *foxc1b* and *acta2* co-expression. These results are consistent with other reports in mouse (Hartmann *et al*., 2015; Grant *et al*., 2017), such as the recently observed zonation of mural cells in the mouse brain (Vanlandewijck *et al*., 2018).

### Transcriptomic differences among embryonic mural cells

We used a combination of two markers, *acta2* (vascular and visceral smooth muscle, in addition to cardiac and some skeletal muscle cells (Whitesell *et al*., 2014)) and *foxc1b* (vascular smooth muscle cells but also pharyngeal arch, ceratohyal, and migratory cranial mesenchymal cells in zebrafish (Jolanta M. Topczewska *et al*., 2001; Miesfeld and Link, 2014)) for FACS to isolate vascular mural cells. This restricts our population to an enriched vascular smooth muscle population. Analyzing the results of the RNA-Seq experiment, it appears that using a double sorted population for both *foxc1b* and *acta2* is a powerful strategy for identifying and isolating vascular smooth muscle cells, as each marker alone is expressed in other cell types, but the intersection of the two markers marks a strongly enriched population of vascular smooth muscle cells.

Within the transcriptomes, we observe strong expression of expected genes such as *acta2, tagln, tpm4b,* and *desmb,* all of which show enriched vascular or visceral smooth muscle expression by *in situ* hybridization. Other muscle related genes (for example, myosins, ion channels, and receptors) are present, and we also identify many genes which are novel, denoted only by chromosomal location that should be investigated as potential mural cell markers. A few genes we tested were not expressed in mural cells; however, we found that most were expressed near blood vessels (data not shown), suggesting the enzymatic dissociation prior to sorting may not have created single cell suspensions (Vanlandewijck *et al*., 2018). Inclusion of a comparison to an endothelial transcriptome (*kdrl:HRAS-mCherry*) reduced the potential for endothelial contamination in the dataset that may arise from incomplete dissociation of cells prior to sorting.

Our smooth muscle cell transcriptomes are not only the first zebrafish mural cell transcriptomes, but also one of the few to profile mural cells in early development. We sorted zebrafish cells at 4 dpf, near the onset of smooth muscle marker expression, whereas most mouse transcriptomic studies used postnatal or adult mice (Lee *et al*., 2015; He *et al*., 2016; Vanlandewijck *et al*., 2018). One study using *pdgfb* and *pdgfrβ* mutant mice at E17.5 (Bondjers *et al*., 2006), was also embryonic, but is at a stage multiple days after the onset of classic mural cell marker expression in embryonic mice (Lindahl *et al*., 1997; Hellström *et al*., 1999). In postnatal mice, *pdgfrβ* and *ng2* double positive pericytes have been sequenced (He *et al*., 2016) and compared to four other pericyte studies, resulting in a pooled brain mural cell transcriptome. The transcriptomes of mouse visceral smooth muscle(Lee *et al*., 2015), and single cell transcriptomes of the brain vasculature to identify endothelial, smooth muscle, pericyte, fibroblast, and astrocyte cells (Vanlandewijck *et al*., 2018) are complementary to our data.

Comparing our smooth muscle transcriptomes (*acta2, foxc1b + acta2*) to the previously published mouse smooth muscle transcriptome datasets (Lee *et al*., 2015; Vanlandewijck *et al*., 2018), we find many genes shared between the datasets, as expected. These include the common mural cell markers *acta2, tagln, desmb, cnn1b, myl9a,* and *mylh11a* that are also observed in colonic and jejunal smooth muscle transcriptomes (Lee *et al*., 2015), and arterial, arteriolar, or venous smooth muscle cell transcriptomes (Vanlandewijck *et al*., 2018). As our *acta2:EGFP* transcriptome will contain transcripts from vascular and visceral smooth muscle, it is expected that we find many common genes. However, *foxc1b:EGFP + acta2:mCherry* represents an enriched vascular smooth muscle cell transcriptome, making it unique. There are 303 genes shared between the *acta2* and *foxc1b + acta2* datasets, and 1644 genes specific to the *foxc1b + acta2* dataset. These include several hundred novel genes that will be interesting to pursue. Common smooth muscle genes are present, including *lmod1, fhl2a, elnb, mylh11a, myocd*, and *kcne4* and have different abundances in the *acta2* and *foxc1b* + *acta2* datasets. For example, *lmod1* is expressed in both datasets, but is more abundant in the *acta2* dataset, and by *in situ* hybridization, *lmod1* shows high expression in visceral smooth muscle. *fhl2a, elnb, myhl11a, myocd, kcne4* are also found in the visceral smooth muscle transcriptomes (Lee *et al*., 2015), and indicate common smooth muscle genes. In addition, there are ∼40 uncharacterized novel genes specific to both the *acta2* and *foxc1b + acta2* datasets, which warrant further investigation as potential smooth muscle specific genes.

In conclusion, we have utilized *foxc1b* expression to identify a window where mesenchymal cells undergo a morphological change to become early smooth muscle cells associated with the endothelium in real-time. This transition requires *foxc1b* for the mesenchymal cells to differentiate into mature, *acta2* expressing smooth muscle cells. In mature smooth muscle, *foxc1b* marks a unique type of smooth muscle, a subset of *acta2* expressing smooth muscle on medium diameter vessels. RNA-Seq data show key gene expression signatures of different subsets of vascular smooth muscle cells. These data will allow for a better understanding of how early smooth muscle cells behave, function, and undergo differentiation.

## Materials and Methods

Complete methods are available in the supplementary methods.

### Ethics, husbandry, and strains

Zebrafish husbandry was performed following standard protocols (Westerfield, 1995). All procedures were approved by the University of Calgary Animal Care Committee, the University of Massachusetts Medical School IACUC, and the University of Alberta Animal Care and Use Committee-Biological Sciences (ACUC-BioSci). Zebrafish strains/transgenic lines include: wild type Tupfel long fin (TL), *Tg(kdrl:mCherry)*^*ci*5^ (Proulx, Lu and Sumanas, 2010), *Tg(kdrl:EGFP)*^*la*116^ (Choi *et al*., 2007), *Tg(kdrl:HRAS-mCherry)*^*s*896^ (Chi *et al*., 2008), *Tg(acta2:EGFP)*^*ca*7^ (Whitesell *et al*., 2014), *Tg(acta2:mCherry)*^*ca*8^ (Whitesell *et al*., 2014), *TgBAC(pdgfrβ:Gal4FF*^*ca*42^ (Ando *et al*., 2016); *UAS-NTR:mCherry*^*c*264^ (Davison *et al*., 2007)), hereafter *Tg(pdgfrβ:mCherry*), *Tg(-5.0kbfoxc1b:EGFP)*^*mw*44^ (Miesfeld and Link, 2014).

### Generation of foxc1a and foxc1b mutant strains

Generation of foxc1a^ua1017^ and foxc1b^ua1018^ CRISPRC mutant strains each required two guide RNAs, produced by cloning into pDR274 (Addgene, #42250), followed by in vitro transcription using the MAXIscript T7 Transcription kit (Ambion, Cat. No. AM1312). *foxc1*^*aua*1017^ target sequences (5’-3’) were AACTCGCTGGGAGTTGTGCC and CCGCCGCCGGAGGGGGGTACACC. *foxc1b*^*ua*1018^ target sequences (5’-3’) were GGCGTTGTGCCTTATATCCC and CGACCGGTGGTGGATATACC.

Cas9 mRNA was synthesized from pMLM3613Cas9 (Addgene, #42251) using a mMESSAGE mMACHINE T7 Transcription kit (Ambion, Cat. No. AM1344), followed by a Poly-A Tailing Kit (Ambion, AM1350). Injections were standardized using 30 ng/µl per gRNA and 300 ng/µl Cas9. Injected fish (P0) were outcrossed and embryos were screened using HRM (Qiagen, Rotor-Gene Q). Mutations were cloned and sequenced from genomic DNA and RNA.

### Sectioning, imaging and image analysis

Confocal images were collected on a Zeiss LSM 700 microscope. Embryos were mounted in 0.8% low melt agarose on glass bottom dishes (MatTek, Ashland, MA, Cat. No. P50G-0-30-F), and were incubated in a heated chamber with the addition of 0.4% Tricaine (Sigma, Cat. No. A5040) for restraint. Representative images are shown.

Adult zebrafish brains (0.5 – 1.5 years old) were dissected and fixed overnight in 4% PFA/1x PBS. Brains were washed in PBS (3x 5 min) before being mounted in a 55°C gelatin solution (15% w/v). Coronal vibratome sections were cut on a Leica Vibratome VT1000S at a thickness between 150 – 300 µm.

Wholemount imaging of stained samples was conducted with a Zeiss Stemi SV11 microscope, with a Zeiss HR camera. Stained wholemount embryos were sectioned at 8 µm after mounting in JB4 resin (Polysciences, Warrington, PA). Sections were imaged on a Zeiss Axio Imager.Z1 microscope with an AxioCam ICc5 camera (Zeiss).

Vessel diameter measurements were measured from confocal images using FIJI/ImageJ (Schindelin *et al*., 2012). Diameters were measured from the external diameter of the endothelium, away from nuclei of mural and endothelial cells. Seven measurements were taken for each sample where possible. Ventral head measurements were taken from the ventral aorta and the aortic arch arteries. In the brain, measurements were predominantly taken from the internal carotid artery (smooth muscle) and central arteries (pericytes). Measurements represent mean vessel diameter ± standard deviation in micrometers.

### *In situ* hybridization and antibody staining

*In situ* hybridization was performed using the method of Lauter *et al*. 2011 (Lauter, Soll and Hauptmann, 2011), with the following modifications. Embryos were permeabilized in 2% H_2_0_2_/methanol for 20 minutes and depigmented in 3% H_2_0_2_/0.5% KOH in water for 10 minutes. Proteinase K permeabilization was at 10 µg /ml for 1 hour for 4 dpf embryos. EGFP was detected with 1:500 mouse αGFP (JL-8, BD Clontech/Takara Bio USA) and the Vectastain ABC Kit (Vector Laboratories, Burlingame CA, USA) using 1:400 αDIG POD Fab fragment (Roche, Cat. No. 11 207 733 910) and FAM (ThermoFisher, Cat. No. C1311) for green fluorescent probe signal, or mouse αGFP (1:500, JL-8, BD Clontech/Takara Bio USA) and Alexa555 goat α mouse fluorescent secondary antibody (1:500, ThermoFisher, Cat. No. A21422). Primers for *in situ* hybridization are listed in Supplemental Table 3.

### FACS & RNA isolation

The protocol for single cell dissociation was based upon Rougeot *et al*. 2014. 300 *acta2:EGFP* and 300 *foxc1b:EGFP* + *acta2:mCherry* (4 dpf) fish were anesthetized with 0.4% Tricaine (Sigma) and pooled. Embryos were treated for 15 minutes with calcium-free Ringers Solution and triturated 15 times. Dissociation Solution was added and triturated 10 times before placing in a 28.5°C water bath with shaking at 60 rpm and periodic trituration for 1 hour. The reaction was stopped, centrifuged and resuspended in Dulbecco’s Phosphate-Buffered Saline (GIBCO by Life Technologies; REF 14040-133), centrifuged and resuspended in fresh Resuspension Solution. The single cell suspension was filtered with 75 µm, followed by 35 µm filters. Cells were then sorted with a BD FACSAria III (BD Bioscience, San Jose, USA) and collected into a collection solution. RNA Isolation of sorted cells was performed using the Trizol method (Ambion by Life Technologies; Carslbad CA, USA; Cat No. 15596026). For analysis of *kdrl:HRAS-mCherry* cells, larvae bearing transgene were anesthetized at 5 dpf, dissociated into single cell suspensions, fixed, and subjected to fluorescence activated cell sorting (FACS) by the University of Massachusetts Medical School Flow Cytometry Core, as described previously (Quillien *et al*., 2017).

### Next Generation Sequencing and bioinformatics

Library preparations used the REPLI-g Single Cell Kit (Cat. No. 150343, Qiagen) for paired end sequencing of 4 dpf samples, using an Illumina NextSeq Platform. 4 dpf samples were: *acta2:EGFP*^+^, *acta2:EGFP*^-^; and *foxc1b:EGFP*^+^ + *acta2:mCherry*^+^, *foxc1b:EGFP*^-^ + *acta2:mCherry*^-^). RNA isolation and library preparation for *kdrl:HRAS-mCherry* positive and negative cell populations was handled as previously reported (Quillien *et al*., 2017).

The sequencing depth for *acta2:EGFP* positive samples averaged 54.8 million reads and 60.9 million reads for the negative population. The sequencing depth for *foxc1b:EGFP* + *acta2:mCherry* double positive samples averaged 59.7 million reads and 61.5 million reads for the double negative population. The sequencing depth for *kdrl:HRAS-mCherry* samples averaged 52.7 million reads for the positive population and 39.1 million reads for the negative population.

Paired-end reads were aligned to 26 chromosomes and 967 primary assembly scaffolds of the zebrafish genome GRCz11, with star_2.5.3a (Dobin *et al*., 2013). Aligned exon fragments with mapping quality higher than 20 were counted toward gene expression with featureCounts_1.5.2 (Liao, Smyth and Shi, 2014). Normalization and differential expression (DE) analysis was performed with DESeq2_1.20.0 (Love, Huber and Anders, 2014). For DE analysis, the original DESeq2 shrinkage estimator was used to estimate log2 fold change (LFC) for each comparison. False discovery rate (FDR < 0.05) and |LFC| > 1 was used as a cut-off to identify significantly enriched genes. Parameters: GTF: GCF_000002035.6_GRCz11_genomic.ucsc.primary.gff. Genome: danRer11.primary.fa

### Smooth muscle quantification

Heterozygote transgenic *acta2:EGFP* fish were imaged, and then FIJI/ImageJ was used to analyze the length of smooth muscle coverage upon the ventral aorta (Schindelin *et al*., 2012). Measurements were recorded from where the bulbus arteriosus merges with the ventral aorta to the distal tip of smooth muscle expression on the ventral aorta or to the bifurcation point of the ventral aorta, whichever was shorter (Isogai, Horiguchi and Weinstein, 2001).

## Acknowledgments

We would like to thank Charlene Watterston, Jasper Greysson-Wong, Nabila Bahrami, and Danielle Blackwell, in addition to the laboratory of Dr. Peng Huang for their helpful comments on the project and paper. We would like to thank the members of the Flow Cytometry core facility and the Alberta Children’s Hospital Research Institute (ACHRI) Genomics Platform at the University of Calgary.

## Data availability

RNA sequencing data is available from the Gene Expression Omnibus (GEO) site (www.ncbi.nlm.nih.gov/geo/) with accession number GSE119718.

